# REFINING THE USE OF SEED BALLS TO SUPPORT SEAGRASS RESTORATION

**DOI:** 10.64898/2026.06.03.729958

**Authors:** Hannah E. Potter, Stijn den Haan, Benjamin L.H. Jones, Emily Yates, Richard K.F. Unsworth

## Abstract

Seagrass ecosystems are experiencing rapid global decline, and current seed-based restoration efforts are frequently hindered by severe early-life bottlenecks such as decapod predation and bioturbation. To bypass these ecological feedbacks, encasing *Zostera marina* seeds in protective clay matrices (seed balls) presents a promising, low-cost intermediate technology for scalable deployment. To refine this approach, a mesocosm experiment was conducted to evaluate the physical integrity, germination success, and early shoot development of *Z. marina* seeds across various clay formulations (fireclay, bentonite, and Kettering loam) and drying treatments. Seed balls were deployed either on the sediment surface or buried, and their performance was evaluated using a hurdle-style generalised additive modelling approach. Results indicated a clear trade-off between structural robustness and biological performance. While naked seeds (70.3% emergence) and buried fireclay balls (67.2% emergence) exhibited the highest overall emergence and post-emergence shoot length, surface-deployed seed balls formulated with higher proportions of bentonite and loam (e.g., 2:4:1 ratio) demonstrated the superior structural integrity necessary for resisting degradation during deployment. However, these denser matrices significantly reduced emergence probabilities and restricted shoot development compared to naked seeds. Formulations lacking loam (1:3) performed the poorest across all biological and structural metrics.

Despite reduced emergence relative to buried treatments, surface-deployed seed balls still achieved high viability, outperforming typical field germination rates. Although further field-based refinement is required to test formulations under dynamic hydrodynamic and bioturbation pressures, seed balls offer a scientifically robust, scalable mechanism to bypass early-stage predation and support widespread seascape recovery.

## Introduction

Seagrasses form one of the world’s most widespread habitats in shallow coastal waters, found on all continents except Antarctica, however seagrass ecosystems are experiencing rapid losses (Cullen-Unsworth and Unsworth, 2016; Unsworth et al., 2022). Current efforts to restore depleted eelgrass ecosystems commonly utilise seed-based methods, but these are hampered by poor in-situ germination success as a result of seed predation, worm burial and other biological parameters [1-3]. Further research to improve current and develop new methods into seagrass restoration is critical to safeguard these keystone ecosystems for the future [4] Predation of seeds by decapods, is a significant cause of seed losses, up to 73% in some studies, attributed to species such as the Common Shore Crab (*Carcinus maenas*) [5, 6]. Across different studies, lugworms and ragworms favourably predated on eelgrass seeds and sprouting shoots, as well as negatively impact shoots through sediment burial and bioturbation [7]. Investigations into preventing predation suggest that burying seeds into the sediment surface, by around 2cm, can prevent predation and promote restoration [8]. Planting eelgrass seeds encased in clay may help prevent early-stage predation during germination and increase the chance of shoot survival [8] In addition to predation pressures, burial of eelgrass seeds and seedlings by lugworms (*A. marina*) and ragworms (*Hediste diversicolor*) have been observed to severely impair eelgrass recovery and greatly limit the success of seed-based restoration methods [2, 7]. Sediment reworking by lugworms causes excessive burial of seeds, inhibiting hypocotyls of germinating seeds. This can result in profound negative impacts on seed-based restoration methods in environments with high lugworm populations [3]. Encasing eelgrass seeds in a clay-structure may minimise excessive burial by increasing the surface area of seeds and protecting seeds from physical damage caused by lugworms and other invertebrates.

Seagrass planting efforts resolve primarily within coastal or estuarine ecosystems that are accessible on foot or by scuba divers. These environments experience a great variation in light availability, water chemical composition, salinity, physical exposure and other ecological factors which may influence seedling settlement and development. This is most prevalent in populated coastal areas where high nutrient input from agriculture, pollution from sewage outflows, increased turbidity from coastal development and sediment deposition and other factors may reduce the suitability of an environment for seedling survival [9]. Previous studies into seagrass restoration have highlighted that even sites deemed suitable from habitat suitability models still experienced low seedling emergence, expected to be a result of bottlenecks to seed germination and retention caused by site conditions like wind fetch, redox depth and sediment type [10, 11].

However, other studies such as van Katwijk, Thorhaug (9) identified that the relationship between seedling survival and population growth in restoration trials was not confounded by other characteristics such as species, method of planting or environmental characteristics. These contradictory studies highlight the need for further research into mechanisms that affect seagrass growth and survival to improve global seed-based restoration efforts [11].

Terrestrially, seed balls [also recognised as ‘seed coatings’] are widely utilised in restoration for overcoming ecological and environmental feedbacks [12]. The seed ball technique has been used since the times of Ancient Egypt to repair farms after annual floodings of the Nile [13], however the use of seed balls in the marine environment is relatively new. This method of propagation is favoured as seed balls are inexpensive to produce, provides favourable seedling habitats and protects seeds from predation [14]. So far there have been limited studies into eelgrass restoration using seed balls, however previous research presented a high seedling survival rate in balls that were buried in the subtidal zone [8]. Although the state of knowledge regarding seed ball restoration methods is limited, previous studies highlight possible success in this area. In their 2022 article Xu, Zhou (8) reported a seedling survival rate ranging from 27 - 47% across respective sites, They concluded that seed balls presented several other advantages compared to traditional methods, such as cheap materials, no associated pollution and easy production [8]. Coals, Coleman (15) observed less promising results from the use of these balls in temperate Australia with the technique been less effective than alternative seeding methods. The use of seed coating is supported in the 2007 article by [16], where coated seed plots experienced a greater seedling density than compared to uncoated seeds [16]. Planting using seed balls may also allow for restoration in more inaccessible areas, such as subtidal areas as balls may be hand-cast from kayaks or boats, allowing them to sink and settle in the sediment.

The present study investigates the germination success of *Zostera marina* (Common eelgrass) seeds in clay balls of different compositions and treatments to determine how seed balls may be utilised in the future for eelgrass restoration. The premise of using seed balls is that the outer clay layer of the seed ball protects the seed from turbulence and predation, allowing for the seed to germinate safely.

## METHODS

A mesocosm experiment was conducted in aquaria at Swansea University to test the effects of different clay ball treatments on the germination and growth of *Zostera marina* seeds. All treatments were housed in a single flow-through system containing UV-treated, sand-filtered seawater at 30 ppt salinity and maintained at 10°C (±1°C). A 12-hour light:dark artificial light cycle was provided throughout the experiment, which ran from 1st December 2023 to 21^st^ February 2024. Water circulation within the system was maintained using a secondary sump pump and chiller. Water changes were carried out weekly. This involved cleaning the aquarium glass, manually removing epiphytic growth from pots and labels, draining half of the main tank and the entire sump, and replenishing the system with fresh UV-treated seawater.

Eelgrass seeds (as reproductive shoots containing spathes) were collected by hand from healthy meadows at Porthdinllaen, North Wales. Sediment was also collected from the site to replicate the natural substrate conditions of the donor meadow. Collected spathes were stored in netted vats containing circulated, oxygenated seawater. As the leaf material degraded, seeds were released and collected between the nets. Once released, seeds were separated from organic matter and small organisms by filtration, then stored in complete darkness in seawater at 50 ppt salinity and 2°C to prevent premature germination and maintain seed quality. Storage conditions and seed health were monitored regularly through water changes and assessments of seed colour, physical integrity, and disease indicators. Degraded seeds, identified by floating, were removed and discarded.

### Clay composites

The materials used in this study were powdered fireclay (F), bentonite (B), and Kettering loam (KL). Fireclay was selected for its prior use in seed-bomb production, non-reactivity, and neutral pH. Bentonite was chosen due to its stable chemical composition, neutral pH, non-oxidising behaviour, and recognition as a low-hazard material with weak cumulative properties. In addition, bentonite is widely used in construction as a swelling and binding agent, and it was expected to improve the structural integrity of submerged clay balls and reduce early degradation. Kettering loam was included because its nutrient-rich soil-clay composition has previously been shown to support eelgrass germination and growth.

### Preliminary trials

Trials were conducted to identify clay mixtures that maintained shape and structural integrity in seawater. A range of ratios containing fireclay (F) and bentonite (B), together with smaller numbers of mixtures incorporating fine sand (FS) and Kettering loam (KL), were tested.

Each material was blended individually to approximately 1 mm grain size using a food blender, sieved, and then hand-mixed to create the desired ratios. Fresh seawater was gradually added until the mixture reached a workable consistency.

Clay balls approximately 20 mm in diameter were formed by hand and placed into labelled pots containing filtered seawater. The balls were left undisturbed for approximately two weeks, during which they were monitored daily for structural integrity and degradation. Integrity was assessed visually using a scale from 1 to 5. A score of 1 represented near-total disintegration with little structural stability, whereas a score of 5 represented minimal degradation with high structural integrity and shape retention. Eelgrass seeds were not included during this stage of testing. Based on these tests, a series of ratios was defined for use in the main experiment (See Table 1).

**Table 1.**
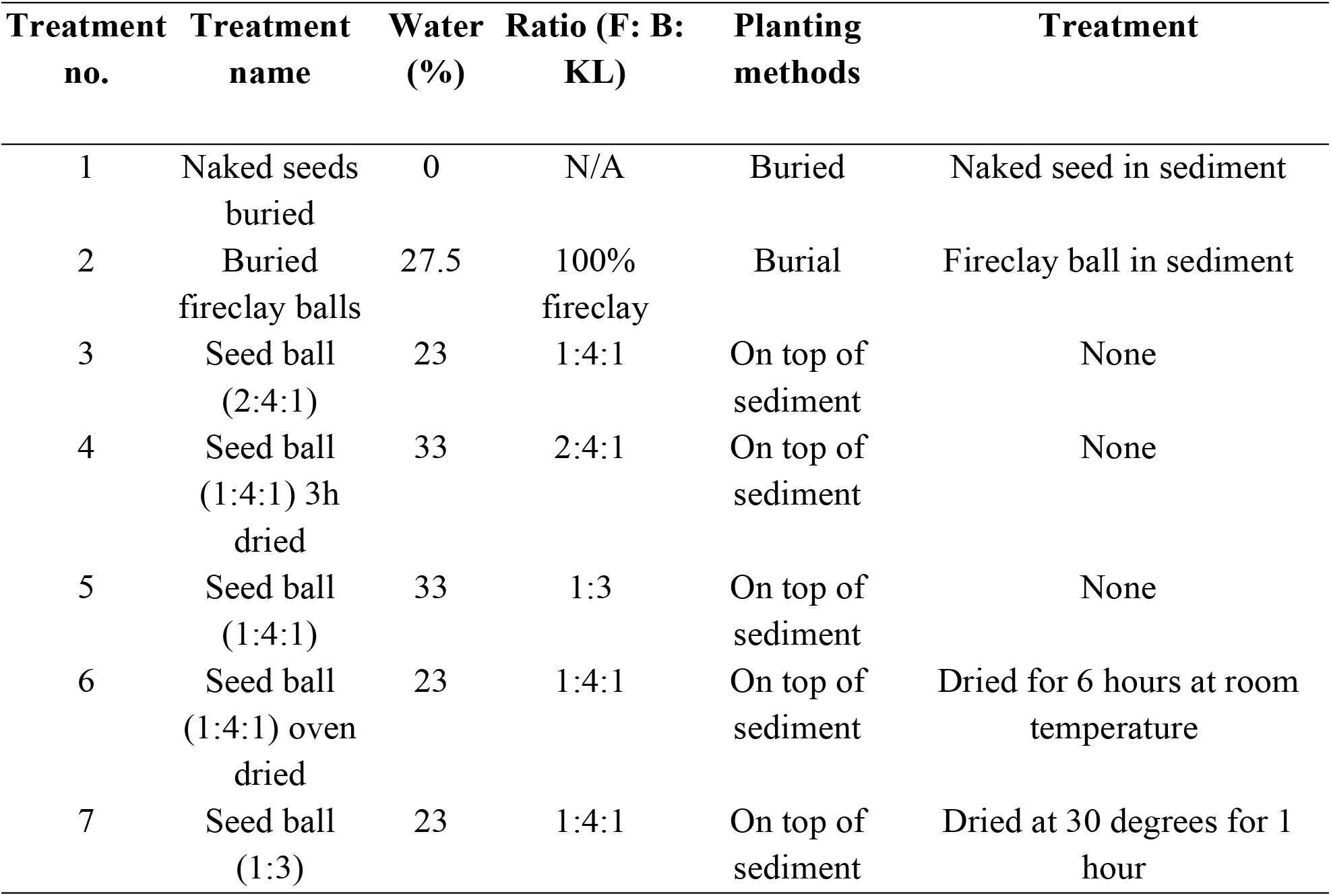
Ratios of clay compositions and water dilutions used in this study, with their respective treatment and planting methods that were used to assess the suitability of each treatment in the germination of Zostera marina seeds.

| Treatment no. | Treatment name | Water (%) | Ratio (F: B: KL) | Planting methods | Treatment |
| --- | --- | --- | --- | --- | --- |
| 1 | Naked seeds buried | 0 | N/A | Buried | Naked seed in sediment |
| 2 | Buried fireclay balls | 27.5 | 100% fireclay | Burial | Fireclay ball in sediment |
| 3 | Seed ball (2:4:1) | 23 | 1:4:1 | On top of sediment | None |
| 4 | Seed ball (1:4:1) 3h dried | 33 | 2:4:1 | On top of sediment | None |
| 5 | Seed ball (1:4:1) | 33 | 1:3 | On top of sediment | None |
| 6 | Seed ball (1:4:1) oven dried | 23 | 1:4:1 | On top of sediment | Dried for 6 hours at room temperature |
| 7 | Seed ball (1:3) | 23 | 1:4:1 | On top of sediment | Dried at 30 degrees for 1 hour |

### Experimental seed ball production

To assess the suitability of clay balls for eelgrass propagation, the selected clay mixtures were produced in larger quantities and formed into seed balls. For each ratio, 64 balls were produced, allowing four replicate trays of 16 balls each. Each tray was 18 x 13 x 8 cm. Seed balls were prepared using the same methods as in the preliminary trials; however, a bait-rolling table was used to produce uniformly sized balls, which were then hand-rolled to a consistent diameter of 18 mm. A single eelgrass seed was inserted into the centre of each ball.

A major concern for the eventual application of these seed balls for field planting is transportation and its impact on drying. Two additional drying treatments were therefore included to examine how drying influenced clay integrity and eelgrass germination success. In the first drying treatment, seed balls (containing the seeds) were air-dried at room temperature for six hours to simulate a short delay between production and deployment. In the second treatment, seed balls were oven-dried at 30°C for one hour to mimic longer-term storage conditions.

All seed balls were placed on sediment layers consisting of play sand and natural sediment collected from Porthdinllaen. Play sand was used as a neutral substrate.

Seed balls from the ‘naked seed’ and the ‘fireclay’ were buried 2 cm into the sediment to assess germination within the substrate. The remaining five treatments were placed on the sediment surface to compare germination success between buried and surface-deployed seed balls. Each planting tray contained two treatments of 16 balls each. Some treatments were deployed immediately after production, whereas others were subjected to drying treatments before deployment.

### Monitoring

Experimental trays were submerged on 1 December 2023, and observations were conducted from 4 January to 21 February 2024. Monitoring was carried out fortnightly to minimise physical disturbance that could affect seedling growth and survival. Weekly water changes were conducted to reduce epiphytic algal growth on trays, in the sediment, and on shoots.

Throughout the experiment, seed ball integrity, seed germination, shoot number and shoot length were monitored visually. Shoot length was measured in millimetres by gently aligning shoots with a ruler using tweezers. Cotyledons were initially included in seedling measurements; however, once seedlings became more developed and produced true leaves, cotyledons were excluded from subsequent observations.

### Data analysis

All data collected in this study is available within the online database Figshare https://doi.org/10.6084/m9.figshare.32328606. We defined seedling emergence as the presence of at least one visible leaf, based on the combined number of cotyledons and green leaves. Cotyledonary leaves were interpreted as newly emerging growth and green leaves as fully developed leaves. Total leaf production was calculated as the sum of cotyledons and green leaves.

Because shoot length could only be measured after visible growth had occurred, we used a hurdle-style modelling approach to separate treatment effects on emergence from treatment effects on subsequent growth [6, 10]. First, using the *mgcv* package for R [17], we modelled the probability of emergence using a binomial generalised additive model (GAM). Second, we modelled total leaf production using a negative binomial GAM, which allowed zero values from non-emerged seeds or seed balls to be retained in the analysis. Third, we modelled shoot length using a Gaussian GAM, restricted to observations where visible growth had occurred. For each model, treatment was included as a fixed effect. Time was modelled using treatment-specific smooth terms to allow growth trajectories to vary among treatments. Smoothness was restricted using a low basis dimension (*k* = 3), reflecting the relatively simple temporal trends expected during early seedling development. Seed or seed ball identity was included as a random effect smooth to account for repeated measurements of the same experimental unit through time. Models were fitted using restricted maximum likelihood.

Next, model predictions were generated for each treatment over the observed experimental period. Predictions were made at the treatment level by excluding seed- or seed-ball-level random effects. For the emergence model, predictions were back-transformed from the logit scale to give probabilities of visible growth. For the leaf count model, predictions were back transformed from the log scale to give predicted leaf counts. For the shoot length model, predictions were generated on the original measurement scale. Modelled responses were visualised with 95% confidence intervals. All analyses were conducted in R [18].

## RESULTS

By the end of the experiment, emergence and early growth differed clearly among treatments (Table 2). Naked seeds had the highest final emergence, with 45 of 64 seeds producing visible growth (70.3%), followed closely by buried fireclay balls, where 43 of 64 seed balls emerged (67.2%). Emergence was lower in all other seed ball treatments, ranging from 50.0% in the 2:4:1 formulation to only 15.6% in the 1:3 formulation. Final leaf production followed the same broad pattern, with naked seeds and buried fireclay balls producing the highest mean number of visible leaves per unit, including non-emerged units as zeros (1.55 ± 1.08 and 1.50 ± 1.13 leaves, respectively). The 1:3 formulation produced the fewest leaves by the end of the experiment (0.30 ± 0.75 leaves).

**Table 2.** Final emergence and early growth across seed ball treatments. Emergence is reported as the number and percentage of seeds or seed balls with visible growth by the final survey. Total leaves are presented as mean ± SD across all units, including non-emerged units as zeros. Shoot length is presented as mean ± SD for emerged seedlings only.

| Treatment | Emergence | Total leaves | Leaf length (mm) |
| --- | --- | --- | --- |
| Naked seeds | 70.3% | 1.55 ± 1.08 | 46.73 ± 12.59 |
| Buried fireclay balls | 67.2% | 1.5 ± 1.13 | 55.65 ± 17.16 |
| Seed ball (2:4:1) | 50% | 1.03 ± 1.1 | 36.19 ± 16.79 |
| Seed ball (1:4:1) 3h dried | 45.3% | 0.78 ± 0.98 | 27.86 ± 18.84 |
| Seed ball (1:4:1) | 42.2% | 0.84 ± 1.09 | 35.33 ± 17.27 |
| Seed ball (1:4:1) oven dried | 40.6% | 0.81 ± 1.1 | 32.96 ± 15.62 |
| Seed ball (1:3) | 15.6% | 0.3 ± 0.75 | 34.7 ± 21 |

Among emerged seedlings, buried fireclay balls produced the longest shoots by the final survey, with a mean shoot length of 55.7 ± 17.2 mm. Naked seeds produced slightly shorter shoots on average (46.7 ± 12.6 mm), while all other seed ball formulations produced shorter shoots, ranging from 27.9 ± 18.8 mm in the 1:4:1 3 h dried treatment to 36.2 ± 16.8 mm in the 2:4:1 treatment.

Seed ball treatment affected early seedling development at multiple stages. Using a hurdle-style modelling approach, we found that treatment influenced not only whether seed balls produced visible growth, but also how many leaves they produced and how long those shoots became after emergence. In all three models, responses changed significantly through time, showing clear temporal development across the experiment.

First, emergence differed significantly among treatments (Fig. 1; R^2^ = 0.943). We found that buried fireclay balls and naked seeds showed the strongest emergence performance, with no detectable difference between them. In contrast, the 1:3 seed ball had a much lower probability of emergence than buried fireclay balls (estimate ± SE = −5.76 ± 1.67, p < 0.001). The 1:4:1 seed ball also leaned towards lower emergence, although this effect was marginal (−2.65 ± 1.49, p = 0.076). The remaining seed ball treatments also showed lower estimated emergence than buried fireclay balls, but these differences were not statistically significant.

**Fig. 1.**
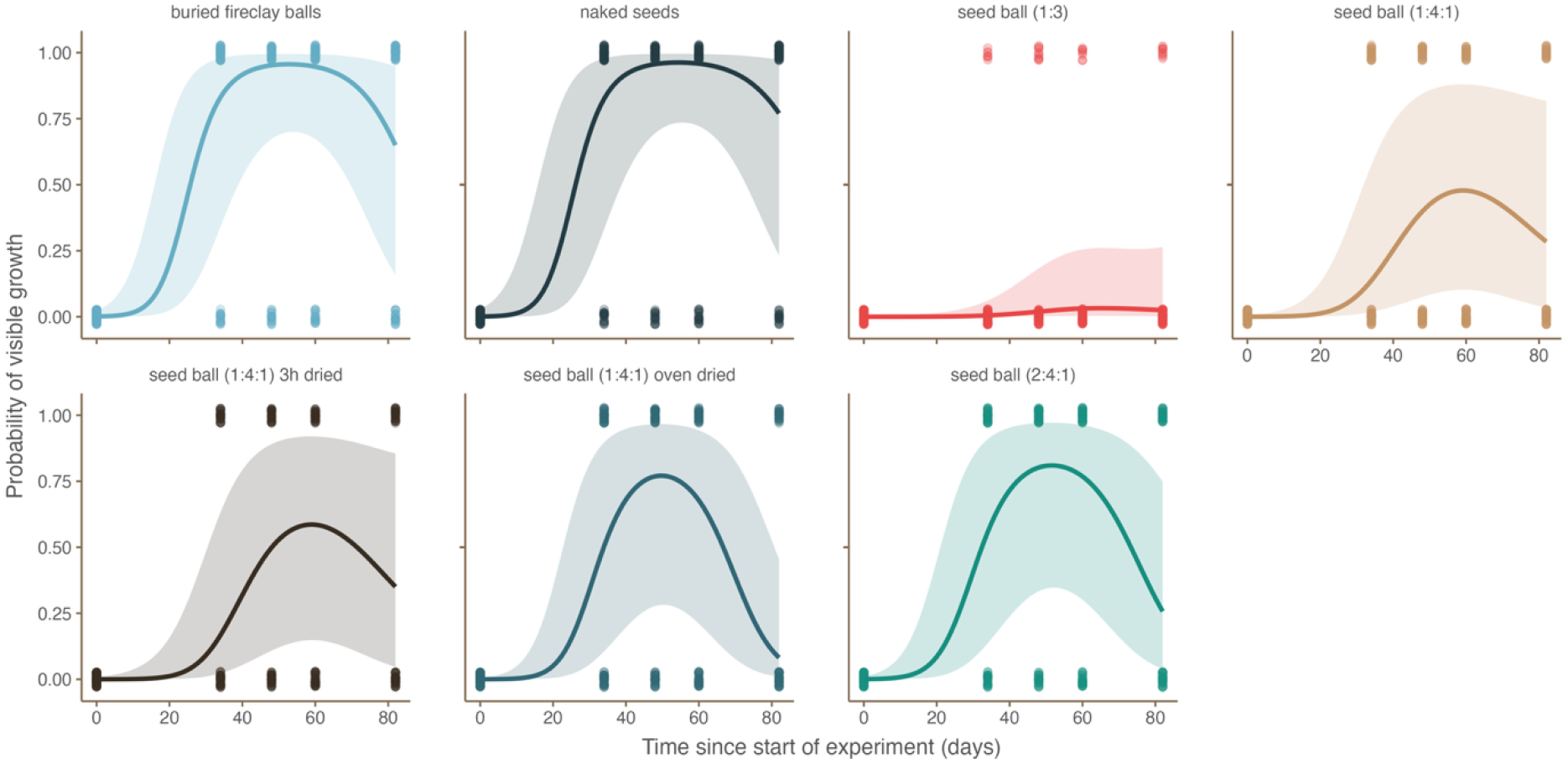
Probability of visible growth across seed ball treatments. Modelled probability of seed ball emergence through time, defined as the presence of at least one visible white or green leaf. Points show observed emergence/non-emergence records, lines show treatment-specific model predictions, and shaded bands show 95% confidence intervals. Predictions are shown at the treatment level, excluding seed ball-level random effects.

Second, leaf production broadly followed a similar pattern. The number of visible leaves increased through time in all treatments (Fig. 2; R^2^ = 0.799). Some seed ball formulations produced fewer leaves than buried fireclay balls. In particular, leaf production was significantly lower in the 1:3 seed ball treatment (−2.11 ± 0.39, p < 0.001), the 1:4:1 treatment (−0.69 ± 0.33, p = 0.037), and the 1:4:1 6h dried treatment (−0.73 ± 0.33, p = 0.029). Naked seeds, the oven-dried 1:4:1 treatment, and the 2:4:1 treatment did not differ significantly from buried fireclay balls in total leaf production.

**Fig. 2.**
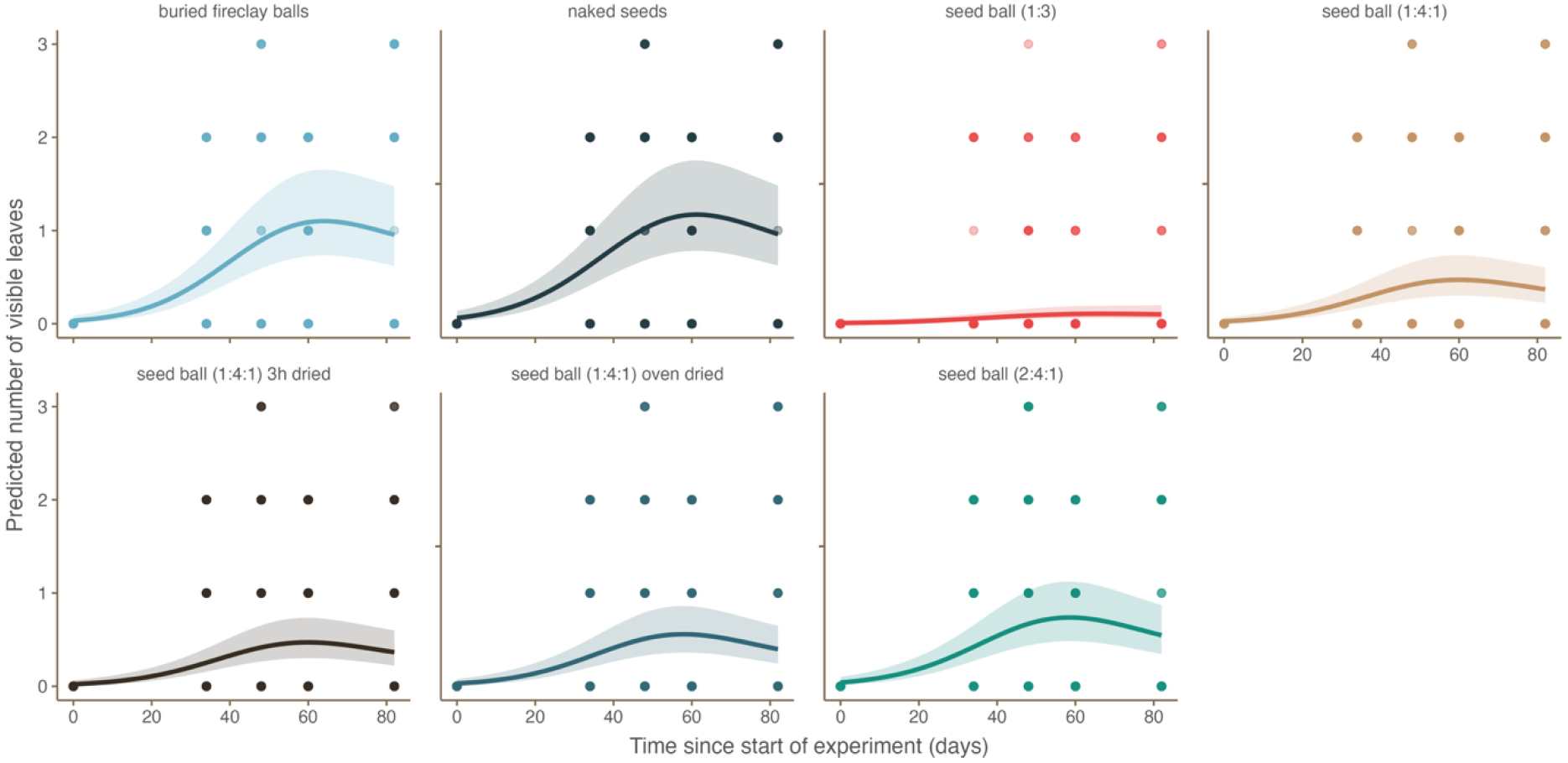
Leaf production across seed ball treatments. Modelled total number of visible leaves through time across seed ball treatments. Total leaves were calculated as the sum of white and green leaves. Points show observed leaf counts, lines show treatment-specific model predictions, and shaded bands show 95% confidence intervals. Predictions are shown at the treatment level, excluding seed ball-level random effects.

Third, post-emergence shoot growth showed a far stronger and more consistent treatment effect. Once visible growth had occurred (i.e., accounting only for balls where leaves had emerged), shoots were shorter in all seed ball formulations than in buried fireclay balls (Fig. 3; R^2^ = 0.683). Shoot length was significantly lower in the 1:3 treatment (−9.01 ± 2.51 mm, p < 0.001), the 1:4:1 treatment (−9.16 ± 1.72 mm, p < 0.001), the 1:4:1 6h dried treatment (−14.13 ± 1.68 mm, p < 0.001), the 1:4:1 oven-dried treatment (−10.76 ± 1.64 mm, p < 0.001), and the 2:4:1 treatment (−9.83 ± 1.58 mm, p < 0.001). Naked seeds again did not differ significantly from buried fireclay balls (−1.38 ± 1.49 mm, p = 0.355).

**Fig. 3.**
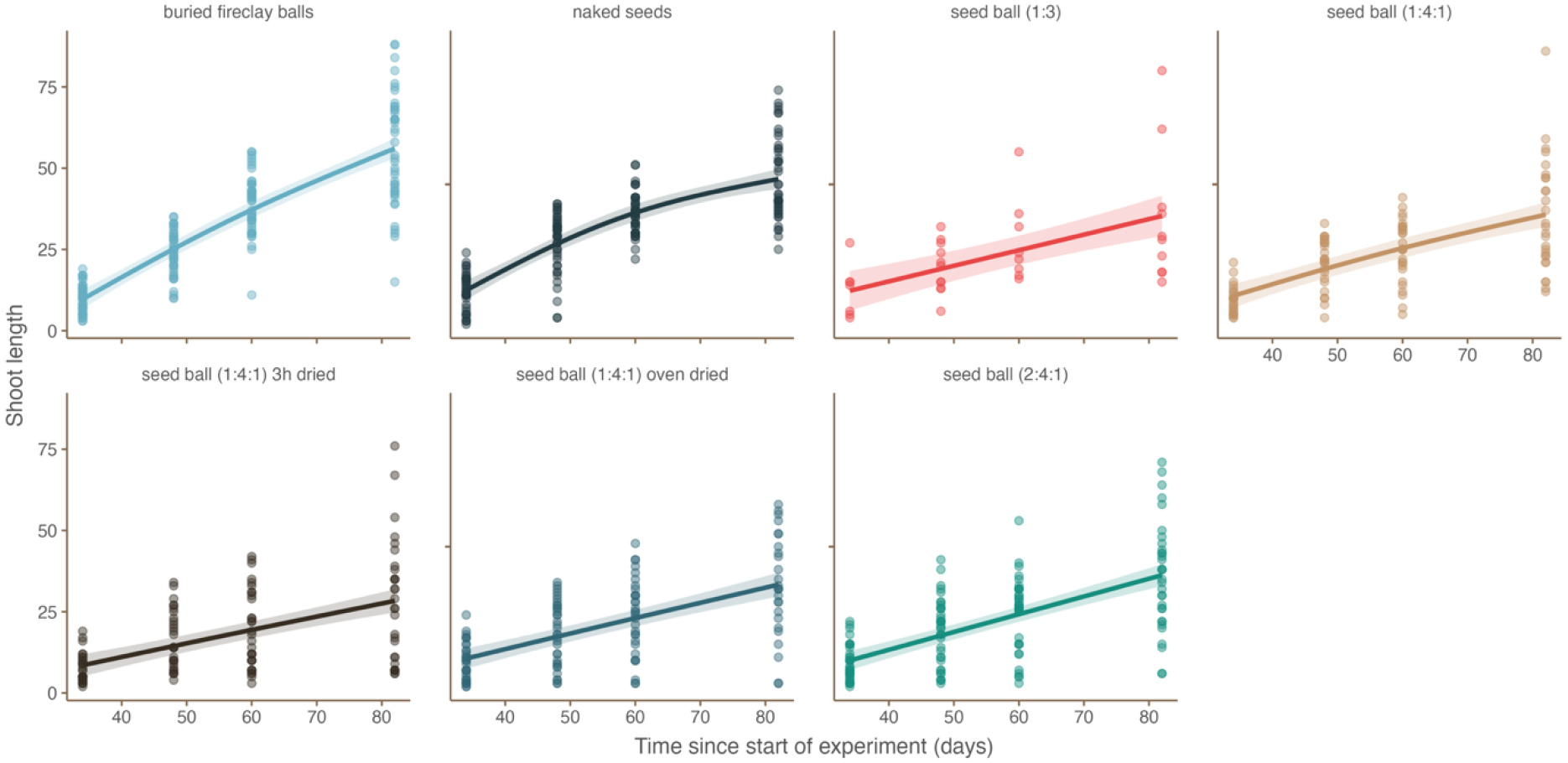
Shoot length across seed ball treatments following emergence. Modelled shoot length through time for seed balls that produced visible growth. Points show observed shoot length measurements, lines show treatment-specific model predictions, and shaded bands show 95% confidence intervals. Predictions are shown at the treatment level, excluding seed ball-level random effects.

Seed ball integrity also varied strongly among formulations and declined significantly through time in all seed ball treatments (Fig. S1). The 1:3 formulation had the lowest mean integrity score, indicating the greatest structural degradation. Notably, the most rapidly degrading formulation also had the weakest biological performance. Therefore, faster degradation did not translate into improved early growth.

Using a hurdle-style modelling approach, we show that seed ball formulation can influence both the likelihood and quality of early seedling development in lab settings. The clearest negative response was observed in the 1:3 formulation, which performed poorly across all three models (emergence, leaf production, and shoot length). The 1:4:1 and dried 1:4:1 formulations also showed reduced performance in some stages of development. By contrast, buried fireclay balls and naked seeds consistently performed best, with naked seeds showing no significant reduction relative to buried fireclay balls across emergence, leaf production, or post-emergence shoot length.

## DISCUSSION

This study provides a novel advancement in the use of seed balls in seed-based seagrass restoration efforts. Across the study, we found strong evidence that seeds germinated across all seed ball mixtures and approaches in a mesocosm setting, paving the way for more targeted application in the field. They were also able to withstand periods of drying, potentially aiding with their use in the field, where transportation and other logistics might delay immediate planting.

Typically, seed germination (and/or emergence) in the field can be lower than 5% [11]. In the present trial, we recorded this at over 40% for all treatments, except one (which didn’t contain Kettering loam), highlighting the viability of these methods. Consistently, burial of seeds has been found to be an optimal means of achieving the highest emergence rates [19, 20]. Here we record both buried treatments (seed ball and naked seed) to have the highest emergence, highlighting the benefits of burial. A potential logistical benefit of using seed balls would be to improve planting efficiency in challenging environments such as soft mudflats or turbid waters. Terrestrial seed planting is increasingly using seed balls and technology to facilitate this [21]. The remote or removed deployment of these balls wouldn’t facilitate seed burial and would likely lead to their placement on top of the sediment. Although our evidence finds a reduced emergence and growth rate from unburied balls relative to buried balls, emergence from some treatments remains very high relative to field-based rates, and leaf growth is good, highlighting the potential of seed ball field deployment methods.

The structural integrity of seed balls is important because their physical characteristics are critical for successful seed dispersal and establishment. Seed balls must be sufficiently firm to withstand transport and aerial dispersal [22], or in a subtidal sense dispersal through the water column. This needs to be ensured while also absorbing moisture on the seabed to facilitate seed imbibition, a process essential for successful germination [23]. In addition, the clay matrix must remain sufficiently porous to facilitate rapid hypocotyl emergence and subsequent leaf development. Here we recorded significant differences in seed ball integrity with respect to formulation. The 1:3 formulation without any loam had the lowest mean integrity score, indicating the greatest structural degradation, less likely to survive being dropped onto the seabed. Notably, the most rapidly degrading formulation also had the weakest biological performance. The dried treatments and the higher-clay formulation (2:4:1) exhibited the best integrity, indicating they may be better formulations for challenging deployments.

The success of treatments with higher clay and loam confirms a range of studies that show the relatively high nutrient content of loam and the properties for absorbing nutrients and developing a complex microbial community [24]. Previous studies on other seagrasses find these sediment types to be effective for lab-based seagrass growth [25]. Fireclays are also now commonly used in various seed injection methods, too, as a medium.[26].

Here we recorded a clear trade-off between the physical integrity of the seed balls and their biological performance. Although formulations with a higher bentonite and loam content, such as the 2:4:1 and dried 1:4:1 treatments, exhibited the best structural integrity for withstanding challenging deployments, this denser composition resulted in reduced seedling emergence and restricted shoot length compared to naked seeds or buried fireclay.

The refinement of these seed ball methods presents an accessible, intermediate technology for marine restoration. By removing the reliance on complex, capital-intensive mechanised deployment or scuba diving, seed balls can be hand-cast from small vessels directly into subtidal environments. This provides a practical means of scaling up restoration efforts and integrating community-led deployment into wider seascape recovery. To build upon these mesocosm results, further research is required in dynamic field settings. Applying these higher-integrity formulations in locations with high hydrodynamic exposure and known bioturbation pressures will test their ability to resist degradation and decapod predation in-situ.

Ultimately, successful seed-based restoration must overcome the significant early-stage bottlenecks of predation and physical displacement [11]. The present study demonstrates that encasing *Zostera marina* seeds in targeted clay formulations can provide the protection required to bypass these ecological feedbacks [3]. While further refinement of the clay matrix is needed to balance integrity with early growth, seed balls offer a scientifically robust, low-cost tool to re-establish functioning eelgrass ecosystems.

---

**Plate 1.**
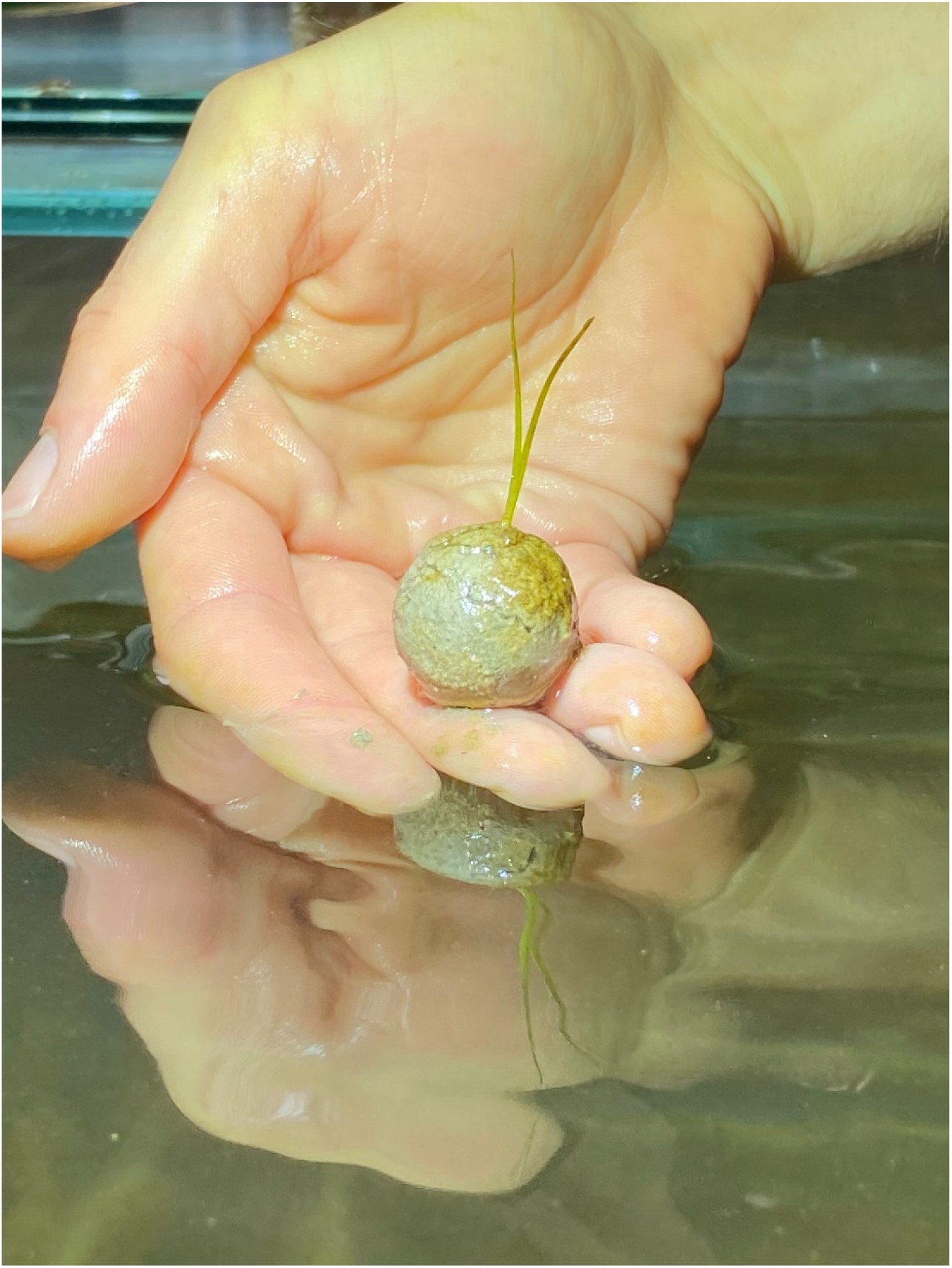
Seagrass seedling emerging from a clay seed ball within the current experiment. Photo: Hannah E. Potter

## Notes

### Competing Interest Statement

The authors have declared no competing interest.

